# Timing and duration of phenological subsidies: toward a mechanistic understanding of impacts on community structure and ecosystem processes in stream food chains

**DOI:** 10.1101/768556

**Authors:** Gaku Takimoto, Takuya Sato

## Abstract

Phenological resources are common across many ecological communities, and can strongly affect community dynamics. Recent field manipulation experiments in stream food chains found that seasonal timing and duration of terrestrial prey inputs affected the feeding behavior, growth, and maturation of fish predators, caused predator-mediated indirect effects on aquatic prey, and modified trophic-cascading effects on litter processing. These experiments described impacts of resource phenological changes over a few month period, and long-term impacts of continued changes in resource phenology are unknown. Here we develop a mathematical model to extrapolate long-term predictions about the effects of changes in resource phenology from the results of field manipulation experiments. The model predicts that advanced timing generally decreases aquatic prey and litter processing and prolonged duration will either increase or decrease aquatic prey and litter processing depending on the total amount and pre-disturbed timing and duration of terrestrial prey inputs. Importantly, our modeling approach clarifies the mechanisms by which stage-specific responses of life history processes in fish, such as growth, maturation, and reproduction, respond to phenological changes in terrestrial prey inputs and mediate indirect effects on aquatic prey and litter processing. Stage-specific responses of life history processes are an integral part of the mechanisms with which to predict the consequences of phenological species interactions at the community and ecosystem levels.

## INTRODUCTION

Many ecological communities occur in the seasonal environment, where the phenology of interacting species determines their relationships (Winemiller and Jepsen 1998; Nakano and Murakami 2001; Power et al. 2008; McMeans et al. 2015). Phenological species interactions generally occur at particular life history stages of interacting species, and affect the physiology, behavior, and demographic rates of organisms at these stages (Post et al. 2008; Seebens et al. 2009; Yang and Rudolf 2010). Phenological interactions at specific life history stages further alter the population dynamics of interacting species (Yang and Rudolf 2010; Visser and Gienapp 2019), and potentially those of other interrelated species in communities and ecosystem processes driven by their activities (Donnelly et al. 2011). With growing evidence of climate change altering species phenology (Post 2018), we need better understanding of whether and what propagating impacts phenological modifications of species interactions can have on community dynamics, structures, and ecosystem processes.

To explore the propagating effects of phenological changes in species interactions in stream food chains, we conducted a series of field experiments (Sato et al. 2016)(Sato et al. in prep). The food chain was comprised of litter detritus as basal resource, aquatic detritivorous arthropods as primary consumers, fish predators preying upon aquatic arthropods as top predators, and terrestrial arthropods supplied from riparian forests to streams as prey subsidies to fish predators (Fig. 1). The fish population was stage-structured into two broad size classes: small and large. Small fish were primarily YOY fish, and large fish included subadult and adult fish. Small fish fed mainly on aquatic prey and secondarily on terrestrial prey subsidies, whereas large fish utilized terrestrial prey subsidies as main diet (Sato and Watanabe 2014). To simulate the inputs of terrestrial prey that occurred seasonally during summer in real streams, we supplied a controlled amount of terrestrial prey subsidies to experimental food chains for a prescribed period. We conducted two experiments. In one experiment, we manipulated the seasonal timing of terrestrial prey inputs; and in the other, we varied the duration of the inputs. Below we briefly explain results from these experiments (summarized in Table 1).

**Table 1.** Empirical observations from our field experiments. Observations I, II, V, and VI are used to incorporate advanced timing and prolonged duration into the model.

|  | Empirical observation | Reference |
| --- | --- | --- |
| I | Advanced timing increased the growth efficiency of small fish feeding on terrestrial prey ($g_S$). | Sato et al. 2016 |
| II | Advanced timing increased the maturation rate of large fish. Accelerated maturation is translated into an increase in the reproduction efficiency of large fish ($r$). | Sato et al. 2016 |
| III | Advanced timing decreased aquatic prey. | Sato et al. 2016 |
| IV | Advanced timing decreased litter processing. | Sato et al. 2016 |
| V | Prolonged duration decreased the proportion of subsidies utilized by small fish ($\omega$) and increases the proportion utilized by large fish ($1 - \omega$) . | Sato et al. in prep. |
| VI | Prolonged duration resulted in small fish consuming a smaller amount of terrestrial prey and a larger amount of aquatic prey, suggesting switching from terrestrial prey foraging ($f$) to aquatic prey foraging ($1 - f$). | Sato et al. in prep. |
| VII | Prolonged duration decreased the growth of small fish, although statistically insignificantly. | Sato et al. in prep. |
| VIII | Prolonged duration increased the growth of large fish and accelerated their maturation. | Sato et al. in prep. |
| IX | Prolonged duration decreased aquatic prey. | Sato et al. in prep. |
| X | Prolonged duration decreased litter processing | Sato et al. in prep. |

**Figure 1.**
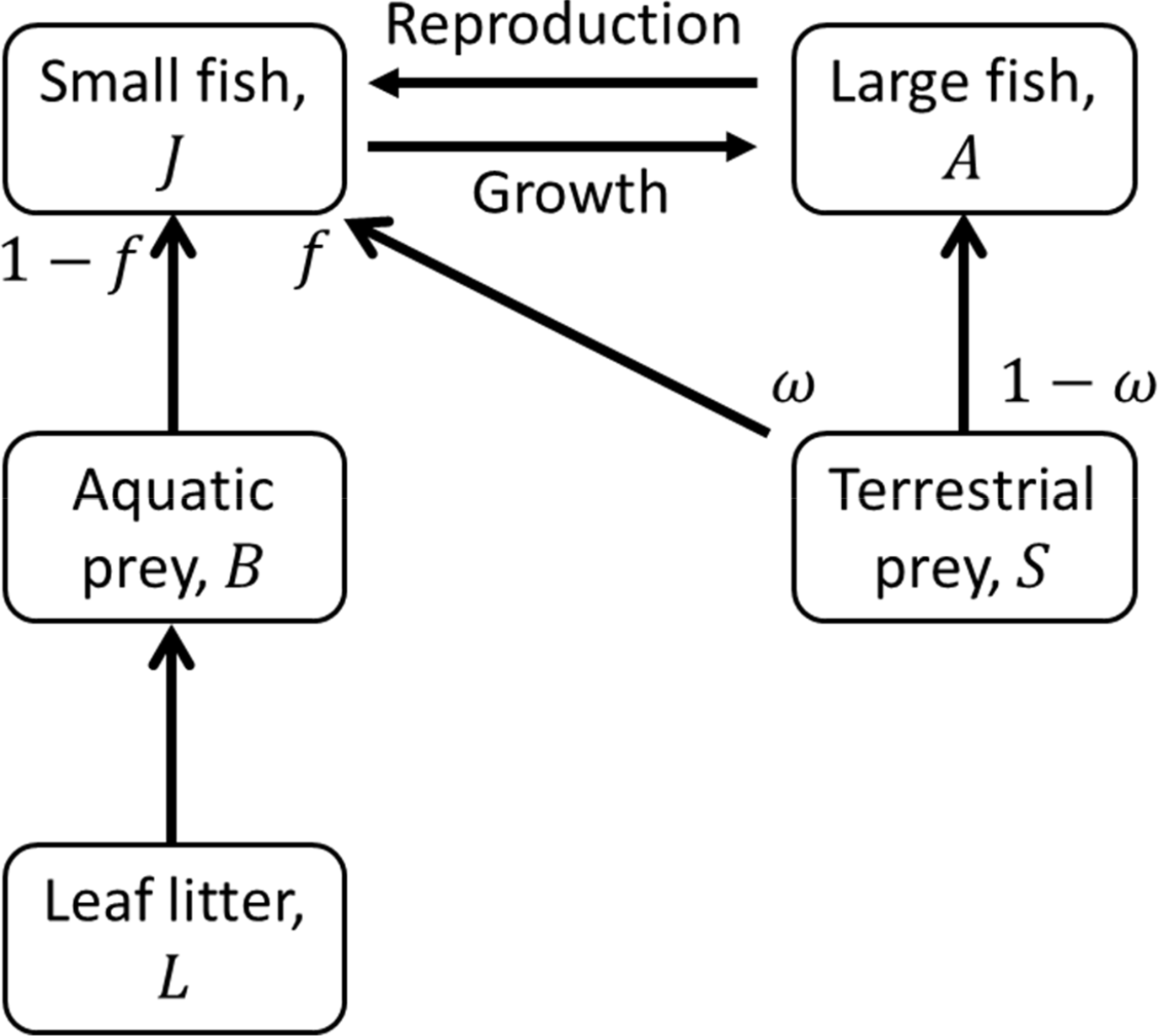
The modelled stream food chain. Longer duration of terrestrial prey subsidy inputs reduces the proportion of subsidies available to small fish (*ω*), and the proportion of time used by small fish to forage subsidies (*f*). Earlier timing of subsidy inputs promotes the reproduction of large fish and the growth of small fish feeding on subsidies.

In the experiment manipulating the timing (Sato et al. 2016), we set up enclosures in natural streams, and stocked a cohort of small fish. We subsidized each enclosure with terrestrial prey early (spring) or late (autumn) in the growing season. We measured the growth rate and the maturation probability of fish, aquatic prey biomass, and the rate of litter processing. We found faster growth and higher maturation probability in fish from enclosures receiving early inputs of terrestrial prey. We also found that the aquatic prey biomass and the litter processing rate were lower when enclosures received early supplies of terrestrial prey.

In the experiment manipulating the duration (Sato et al. in prep.), we established stream food chains in mesocosms. We considered two duration levels: pulsed and prolonged. We kept the total input amount supplied to each mesocosm constant at a realistic level. We measured the individual growth of fish, the consumption of aquatic and terrestrial prey by fish, the stage structure of fish, aquatic prey abundance, and the rate of litter processing. We found that large fish fed on a larger amount of terrestrial prey, grew better, and reached maturity at a higher rate in the prolonged than in the pulsed treatments. Small fish, on the other hand, fed less on terrestrial prey and grew more slowly in the prolonged treatment. Aquatic prey abundance and litter processing rate at the end of the experiment were smaller in the prolonged treatment.

Our field experiments demonstrated that phenological changes in species interactions affected life history processes of fish, such as growth and maturation, and caused indirect impacts on community structure and ecosystem processes. However, our experiments have two weak points, although common to many field experiments in general. First, our experiments lasted for a single growing season, although phenological modifications under climate change will persist for long. It remains unknown to what extent the results of the short-term experiments could inform long-term effects of persistent phenological changes. Second, our experiments manipulated either duration or timing of phenological subsidy inputs, although on-going climate change may affect both simultaneously (CaraDonna et al. 2014; Post 2018). Our experiments may be limited to infer the double impacts of changing timing and duration. An alternative approach using mathematical models might complement these field experiments to inform longer-term consequences across more various situations.

Here we develop a mathematical model to describe the long-term dynamics of stream food chains studied in our experiments. Incorporating empirical observations gained from our field experiments (Table 1), we aim at generating long-term predictions about the effects of phenological changes in the timing and duration of terrestrial prey inputs on aquatic prey and litter processing, and clarifying the mechanisms and the conditions underlying these effects. Although there have been a number of mathematical models on phenological species interactions (Cushing 1990; Durant et al. 2005; Nakazawa and Doi 2012; Revilla et al. 2014; Rudolf 2019), few have considered communities involving more than three species with an explicit formulation of life history stages at which phenological species interactions occur. By contrast, our model considers five interacting members of stream food chains including two life-history stages of predators. We have previously used mathematical models to study the effects of phenological timing and duration of subsidy inputs on the stability and structure of recipient communities (Takimoto et al. 2002, 2009). These models do not consider specific life history stages at which phenological species interactions occur. Our current model, by formulating stage-specific responses to phenological species interactions, clarifies the mechanisms by which changes in the timing and duration of subsidy inputs cause cascading impacts on community structure and ecosystem processes.

## MODEL DEVELOPMENT

We develop a differential equation model that describes the dynamics of a stream food chain subsidized with terrestrial prey (Fig. 1). We incorporate the results of our filed experiments (Table 1) into this model. The model has four state variables: leaf litter abundance (*L*), aquatic prey abundance (*B*), the abundance of small (juvenile) fish predators (*J*), and the abundance of large (subadult and adult) fish predators (*A*). The following model equations are used:

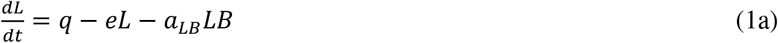

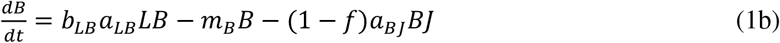

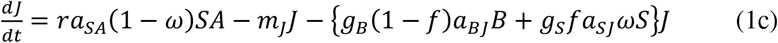

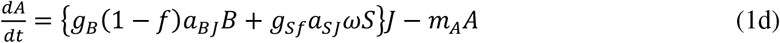

The parameter *S* denotes the availability of terrestrial prey subsidies to fish predators. We consider that the availability is controlled by extrinsic factors of donor terrestrial ecosystems. Other symbols used in the equations are listed and defined in Table 2. This model describes food-chain dynamics on the time scale of years. That is, values of state variables and parameters are their annual averages, and the model follows the changes of these annual averages through multiple years. This means that the duration and timing of subsidy inputs are not explicitly parameterized in the model. Rather, we express changes in the duration and timing of subsidy inputs as the changes of model parameters.

**Table 2.** Symbols and definitions of state variables and parameters in the model, and default parameter values used in numerical analysis.

| Symbol | Definition | Default value |
| --- | --- | --- |
| $L$ | Litter abundance | |
| $B$ | Aquatic prey abundance | |
| $J$ | Small fish abundance | |
| $A$ | Large fish abundance | |
| $q$ | Input rate of litter | 10 |
| $e$ | Loss rate of litter | 0.1 |
| $a_{LB}$ | Consumption coefficient of litter by aquatic prey | 1 |
| $a_{BJ}$ | Consumption coefficient of aquatic prey by small fish | 1 |
| $a_{SJ}$ | Consumption coefficient of terrestrial prey by small fish | 1 |
| $a_{SA}$ | Consumption coefficient of terrestrial prey by large fish | 1 |
| $b_{LB}$ | Conversion efficiency of aquatic prey consuming litter | 0.2 |
| $m_B$ | Mortality of aquatic prey | 0.5 |
| $m_J$ | Mortality of small fish | 0.3 |
| $m_A$ | Mortality of large fish | 0.3 |
| $g_B$ | Growth efficiency of small fish feeding on aquatic prey | 0.1 |
| $g_S$ | Growth efficiency of small fish feeding on terrestrial prey | 0.1 |
| $r$ | Reproduction efficiency of large fish | 0.1 |
| $S$ | Total inputs of terrestrial prey | 10 |
| $\omega$ | Proportion of terrestrial prey available to small fish | 0.2 |
| $f$ | Proportion of time used by small fish to forage terrestrial prey | 0.2 |

**Table 3.** Responses of fish population to phenological changes in terrestrial prey inputs

|  | Prolonged duration | Advanced timing |
| --- | --- | --- |
| Small fish, $\bar{J}$ | +/- | + |
| Large fish, $\bar{A}$ | +/- | +/- |
| Abundance ratio, $\bar{A}/\bar{J}$ | - | - |

We use parameters *g*_*S*_, *r*, *ω*, and *f* to express changes in the timing and duration of subsidy inputs. Earlier timing is expressed as the increase of *g*_*S*_ and *r*. Empirical observations (I and II in Table 1) suggests that subsidy-derived energy contributes to the growth and maturation of fish more effectively when supplied at earlier timing. We incorporate this observation as the increase of *g*_*S*_ and *r* in response to advanced timing. Since large fish in our model include subadult and adult, we consider that advanced timing accelerates the maturation of subadult and thus contributes to the increase of *r*.

Prolonged duration is expressed as the decrease of *ω* and *f*. Prolonged duration decreases the proportion of subsidies available to small fish (*ω*) and increases the proportion available to large fish (1 − *ω*) (empirical observation V, Table 1). This is because prolonged duration decreases the daily inputs of terrestrial prey and low daily inputs intensify interference by large fish that prevents small fish from accessing terrestrial prey (Sato and Watanabe 2014). In addition, we consider that a reduced proportion of subsidies available to small fish (low *ω*) causes a functional response by small fish that reduces the proportion of time for small fish foraging on terrestrial prey (*f*) (empirical observation VI, Table 1). This means that we assume a positive correlation between *ω* and *f*.

## MODEL ANLYSIS and RESULTS

### Functional and numerical responses by fish

Changes in the duration and timing of terrestrial prey inputs affect the functional and numerical responses of small and large fish, which then alters population and community structures and ecosystem processes. We first illustrate how phenological changes in terrestrial prey inputs affect the functional and numerical responses of fish. Functional response by small fish to aquatic prey is expressed by the term

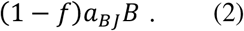

This term shows that prolonged duration (reduced *f*) increases the time spent by small fish to forage aquatic prey and thus increases consumption on aquatic prey, and altered timing has no effect.

Functional response by small fish to terrestrial prey corresponds to the term

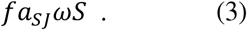

This term indicates that prolonged duration (reduced *ω* and *f*) decreases the availability of terrestrial prey to small fish and also that small fish reduces the time spent to forage terrestrial prey, both of which decrease the consumption of terrestrial prey by small fish. The change of timing has no influence on this term. The responses of these terms (2) and (3) to prolonged duration correspond to empirical observations V and VI (Table 1), which is obvious since we developed the model as such.

Numerical responses of fish to aquatic and terrestrial prey have three components corresponding to the growth of small fish feeding on aquatic prey, the growth of small fish feeding on terrestrial prey, and the reproduction of large fish. The growth of small fish feeding on aquatic prey is represented by the term

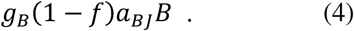

This term means that prolonged duration increases the growth of small fish feeding on aquatic prey as a result of increased consumption of aquatic prey (Expression 2; Fig. 2). There is no effect of changing timing on this growth term.

**Figure 2.**
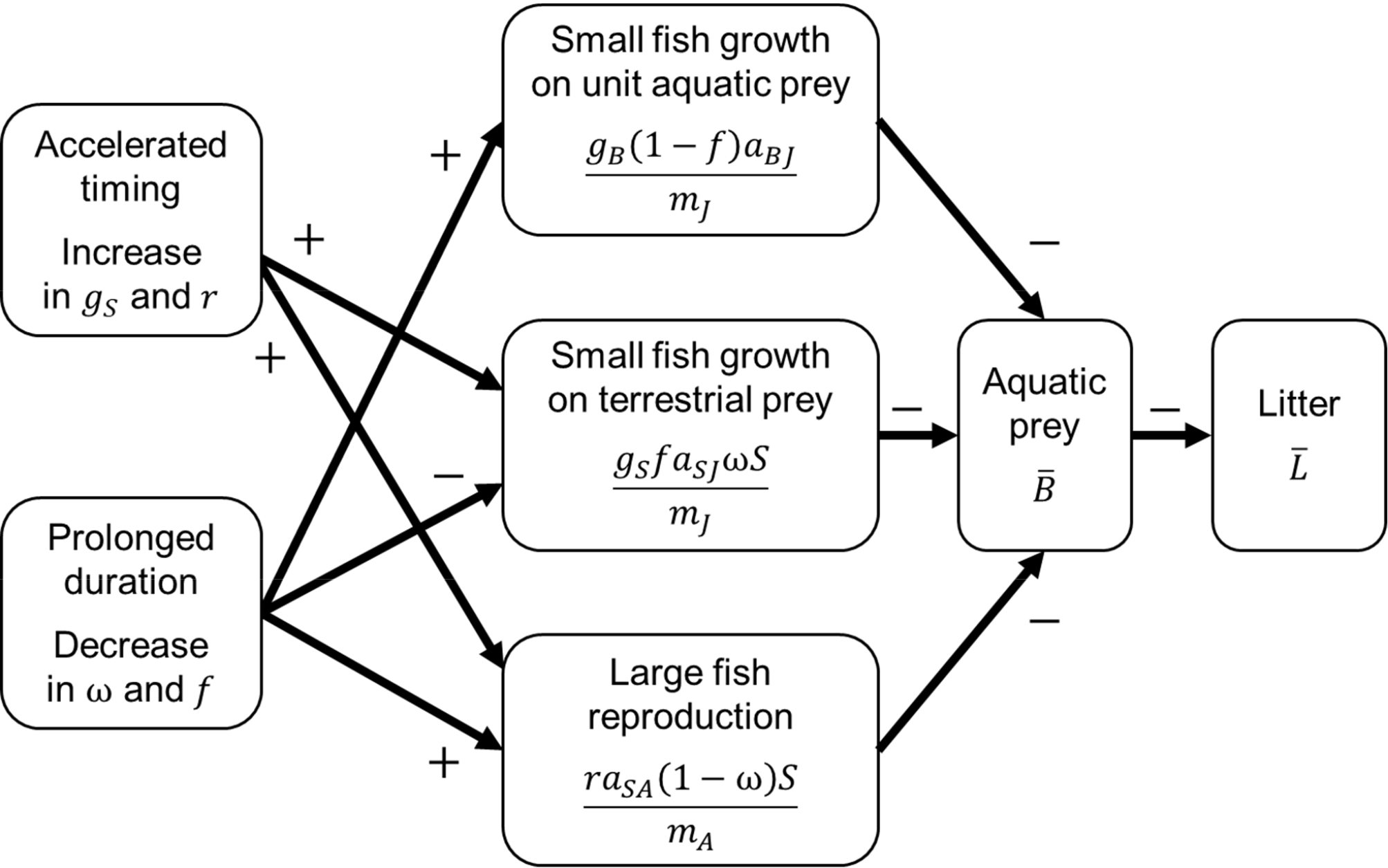
Mechanisms of the responses of life history processes to advanced timing and prolonged duration that mediate their indirect effects on aquatic prey and litter abundance. The plus and minus signs indicate the direction of effects.

The growth of small fish feeding on terrestrial prey is expressed by the term

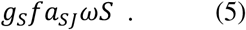

This term shows that prolonged duration decreases the growth, and advanced timing (increased *g*_*S*_) promotes the growth (Fig. 2). The effect of prolonged duration is derived from reduced consumption of terrestrial prey (Expression 3), whereas the effect of advanced timing is due to enhanced efficiency in the conversion of terrestrial prey consumption into growth.

The term expressing the reproduction of large fish is

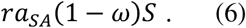

This term shows that prolonged duration and advanced timing (increased *r*) both promote the reproduction of large fish (Fig. 2). The effect of prolonged duration is derived from the increased availability of terrestrial prey to large fish, whereas the effect of advanced timing is due to accelerated maturation.

### Fish population structure

We study the long-term effects of prolonged duration and advanced timing on the abundance of litter, aquatic prey, and small and large fish by examining their equilibrium abundances. At equilibrium where all members in the food chain persist (i.e., have positive abundances), their abundances can be expressed as

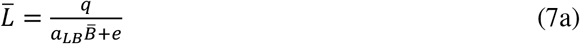

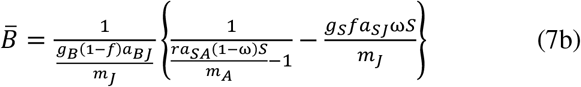

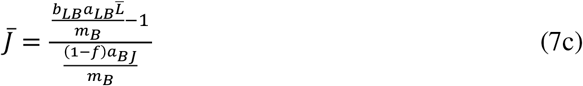

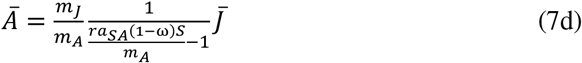

where bars over the variables indicate their equilibrium values. Note that the aquatic prey abundance is expressed as a unique combination of model parameters, whereas the abundances of litter and fish are expressed as functions of model variables in addition to model parameters. Using these expressions, we examine how fish, aquatic prey, and litter processing respond to changes in the duration and timing of terrestrial prey inputs.

Eq. (7c) indicates that the small fish abundance depends positively on the intrinsic production of aquatic prey,

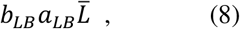

and negatively on the per capita exploitation of aquatic prey by small fish,

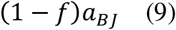

(both are multiplied with the expected life time of aquatic prey, 1/*m*_*B*_, in Eq. 7c). It is difficult to predict how prolonged duration affects the small fish abundance because prolonged duration can either increase or decrease the litter abundance (see below). However, when prolonged duration increases aquatic prey (and litter processing) and decreases the litter abundance, prolonged duration will decrease the small fish abundance because it increases the exploitation of aquatic prey by small fish. On the other hand, advanced timing will uniquely increase small fish because it decreases aquatic prey and increases litter (see below).

Eq. (7c) indicates that the large fish abundance is linearly related to the small fish abundance with a coefficient

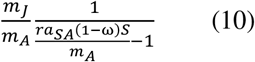

that defines the stage structure of fish population (the abundance ratio of large to small fish). Noting that this abundance ratio depends on the reproduction of large fish in the model term (6), we find that prolonged duration and advanced timing both decrease the ratio of large to small fish abundances by increasing the recruitment of small fish. The absolute abundance of large fish, on the other hand, will show more complex responses to prolonged duration and advanced timing, because the responses of the small fish abundance and the abundance ratio interactively will affect the large fish abundance.

### Aquatic prey abundance

Eq. (7b) shows that the growth and reproduction of fish systematically determines the aquatic prey abundance (Fig. 2). Specifically, the growth of small fish feeding on aquatic prey has a negative effect on aquatic prey through the term

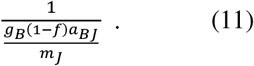

On the other hand, a negative effect of the growth of small fish feeding on terrestrial prey is expressed by the term

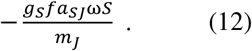

In addition, the term

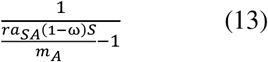

represents a negative effect of reproduction by large fish. These negative effect terms increase in magnitude with the increase of total subsidy input *S*. This shows that numerical response by fish to the increase of overall subsidy inputs through growth and reproduction causes a long-term negative effect on the aquatic prey abundance.

Prolonged duration and advanced timing affect the aquatic prey abundance through modifying these growth and reproduction terms. Compared to the effects of prolonged duration (see below), the effects of advanced timing on the aquatic prey abundance are straightforward (Fig. 2). Advanced timing, expressed as the increase of the growth efficiency *g*_*S*_, decreases the aquatic prey abundance (Eq. 7b) through an increased growth of small fish feeding on terrestrial prey (Expression 12). Similarly, advanced timing, expressed as the increase of the reproduction efficiency *r*, decreases the aquatic prey abundance through the increase of reproduction by large fish (Expression 13). We thus expect that advanced timing generally decreases the aquatic prey abundance.

In contrast, the effects of prolonged duration are more complex (Fig. 2). Prolonged duration may increase the reproduction of large fish (Expression 13), through which a negative effect is transmitted on aquatic prey, but decrease the growth of small fish feeding on terrestrial prey (Expression 12), through which a positive effect will result. Moreover, prolonged duration increases the time spent by small fish to feed on aquatic prey (Expression 11), which mediates a negative effect. Relative strength of these component effects determines the net effect of prolonged duration on aquatic prey.

To explore the effects of prolonged duration, we begin with a limiting case, in which duration is long enough that large fish consume most terrestrial prey without leaving much to small fish. A formal assumption that *ω* ≈ 0 and *f* ≈ 0 corresponds to this case. Under this assumption, a Taylor expansion of 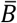 in terms of *ω* and *f* at *ω* = *f* = 0 shows that the aquatic prey abundance is approximated as:

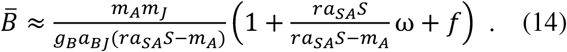

This expression means that when duration is sufficiently long, a further increase of duration (reduced *ω* and *f*) will reduce the aquatic prey abundance. In other words, shortening an originally long duration would increase aquatic prey. This outcome occurs because the growth term of small fish feeding on terrestrial prey becomes effectively zero when *ω* ≈ 0 and *f* ≈ 0 and receive a negligible effect from the change of duration.

To explore more general cases with larger *ω* and *f*, we resort to numerical approach. We assume a simple relationship

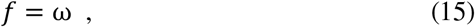

to incorporate their likely positive correlation (empirical observation II). We evaluate the response of aquatic prey abundance to prolonged duration by examining the sign of 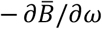.

Figure 3 illustrates that the effects of prolonged duration on the aquatic prey abundance depend on model parameters. For example, the original duration itself affects whether a further increase of duration increases or decreases aquatic prey (Fig. 3A). When subsidy input *S* is low, prolonged duration generally decreases aquatic prey because prolonged duration induces a numerical response by fish to terrestrial prey through enhancing the reproduction of large fish. When *S* is large, on the other hand, prolonged duration rather increases aquatic prey when the original duration is relatively short. When the duration is short, terrestrial prey inputs contribute to the increase of fish through enhancing the growth of small fish feeding on terrestrial prey. Lengthening the originally short duration reduces the growth of small fish feeding on terrestrial prey and weakens the numerical response by fish, causing a positive indirect effect on aquatic prey.

**Figure 3.**
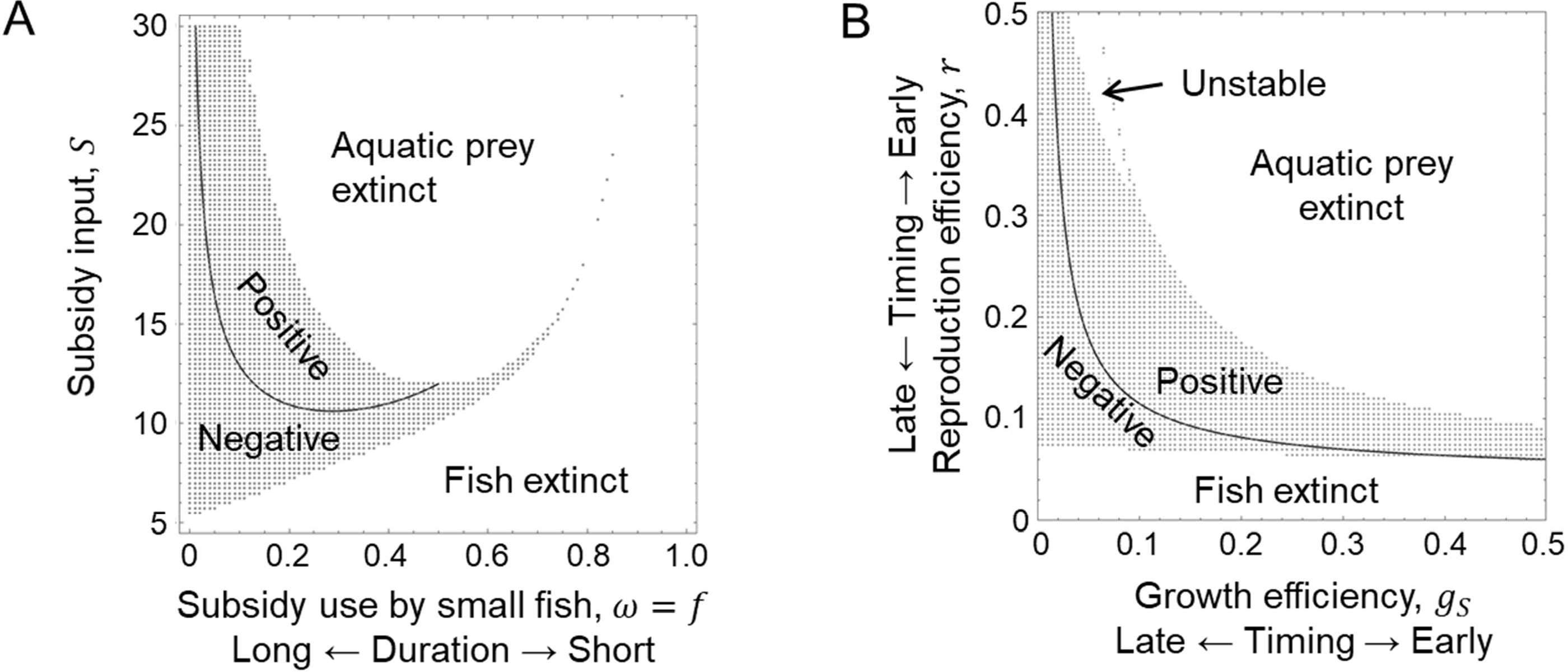
The sign of long-term indirect effect on aquatic prey and litter processing caused by prolonged duration. The hatched areas are parameter regions that yield a locally-stable equilibrium, where large and small fish, aquatic prey, and litter have positive abundances. A, effects of the pre-disturbed proportion of terrestrial prey available to small fish (*ω*) and subsidy input (*S*). B, effects of the growth rate of small fish feeding on subsidies (*g*_*S*_) and the reproduction rate of large fish (*r*). The proportion of time used by small fish to forage subsidies is linearly related to the proportion of subsidies available to small fish (*f* = *ω*). Unchanged parameters are set to their default values listed in Table 2.

The effects of prolonged duration on aquatic prey depend also on other parameters, such as the growth efficiency *g*_*S*_ of small fish feeding on terrestrial prey and the reproduction efficiency *r* of large fish. Noting that *g*_*S*_ and *r* are parameters controlled by the timing of terrestrial prey inputs, we find that the effect of prolonged duration on aquatic prey depends on the timing of terrestrial prey inputs (Fig. 3B). When inputs occur late, fish population suffers from low growth efficiency of small fish (*g*_*S*_) and low reproduction efficiency of large fish (*r*; too low *r* causes fish extinction in Fig. 3B), which allows the proliferation of aquatic prey. Under such situations, prolonged duration that diverts more terrestrial prey to large fish will increase the reproduction of large fish and save fish population, which acts to reduce aquatic prey. When inputs occur early, on the other hand, high efficiency in growth and reproduction of fish utilizing terrestrial prey enhances fish population and aquatic prey in turn suffers from high predation (too large *g*_*S*_ and *r* cause aquatic prey extinction in Fig. 3B). In such a case, reduction of terrestrial prey availability to small fish, a result of prolonged duration, will suppress fish population and save aquatic prey from fish predation to cause a positive effect.

Other parameters, such as the mortality of small and large fish and their consumption efficiencies, should also affect the effects of prolonged duration on aquatic prey. These parameters modulate the effect of prolonged duration by changing the relative magnitudes of component effects through small fish growth and large fish reproduction (cf. Fig. 2).

### Litter processing

The rate of litter processing increases with the aquatic prey abundance, resulting in a negative relationship between the aquatic prey abundance and the litter abundance (Eq. 7a). We thus predict that the effects of prolonged duration and advanced timing on the litter abundance occur in the opposite direction to that of the effects on the aquatic prey abundance. Parameters determining the input and loss of litter and the litter consumption efficiency of aquatic prey will regulate the size of the effects of prolonged duration and advanced timing. For example, processing by aquatic prey will have little impact on the litter abundance and thus the effects of prolonged duration and advanced timing will be small, if the input and loss rates of litter are large.

## DISCUSSION

Many food webs occur in the seasonal environment, in which the phenology of predators and prey defines their interactions (Winemiller and Jepsen 1998; Nakano and Murakami 2001; Power et al. 2008; McMeans et al. 2015). We developed and analyzed a mathematical model that describes the dynamics of stream food chains in which seasonal inputs of terrestrial prey subsidize fish predators. Our model is based on our previous field manipulation experiments that studied the direct and indirect effects of the timing and duration of seasonal terrestrial prey on the behavior and growth of fish predators, the abundance of aquatic prey, and leaf litter processing over a relatively short term (within a year). Our model complements the results of these experiments by generating predictions on the long-term effects of the timing and duration of seasonal terrestrial prey subsidies, as well as the interacting effects of timing and duration. Our approach combining field experiments and mathematical modeling demonstrates that modifications in phenological species interactions change the behavior and life-history processes of interacting species, which in turn causes indirect effects on other interrelated species in communities and ecosystem processes.

Our model employed a number of assumptions that sacrifice realistic details for the sake of simplicity. We discuss potential outcomes of relaxing these assumptions. We have assumed that *ω* and *f* vary in response to changes in the duration of terrestrial prey inputs. In reality, *ω* and *f* can depend also on total inputs *S*. Larger values of *S* increase the availability of terrestrial prey to fish, which might increase *ω* and *f* for a given length of duration. Such an effect could counter the effect of prolonged duration that decreases *ω* and *f*. As a result, the effect of increasing *S* in Fig. 2A could be depicted by an upper-right shift of parameter values on the *ω*-*S* plane instead of a straight upward shift, most likely promoting a positive effect of prolonged duration on aquatic prey and litter processing.

Our model assumed that large fish did not utilize aquatic prey, although in reality large fish could feed on aquatic prey albeit to a lesser extent than terrestrial prey (Nakano et al. 1999; Sato and Watanabe 2014). Incorporating this trophic link into the model would preclude simple expressions of equilibrium abundances. However, we expect that utilization of aquatic prey by large fish (Nakano et al. 1999) might promote the positive effect of prolonged duration on aquatic prey because prolonged duration would induce a functional response by large fish to feed less on aquatic prey.

Our model is focused on litter as the base of stream food chains. Algal production is another important resource base, and our previous experiments looked into the responses of algal abundance to advanced timing and prolonged duration (Sato et al. 2016; in prep). We expect that a similar model should apply to algal-production-based stream food chains, and analogous predictions could be derived for the indirect effects of advanced timing and prolonged duration on aquatic arthropods utilizing algae and algal production.

In our model, life history processes, such as growth and reproduction, of fish responded to advanced timing and prolonged duration (Fig. 2). These responses in the model are in close agreement with empirical observations from our field experiment. Advanced timing was incorporated directly into small fish growth efficiency (*g*_*S*_) and large fish reproduction efficiency (*r*). Thus, close matching between the model and empirical observations (I, II in Table 1) is of no surprise. Prolonged duration in the model decreased the growth term of small fish feeding on terrestrial prey (Eq. 5, Fig. 2). This matches prolonged duration reducing small fish growth in our experiment (although statistically insignificant, VII in Table 1). Also, prolonged duration in our model enhanced the reproduction of large fish (Eq. 6, Fig. 2). This parallels prolonged duration potentially promoting the reproductive maturation of large fish in our experiments (VIII in Table 1).

Long-term predictions from our model complement the relatively short-term results of our field experiments. One of our experiments found that early inputs of terrestrial prey decreased aquatic prey and decelerated litter processing after four months (III and IV, Table 1). The direction of these indirect effects are consistent with the long-term predictions of our model. Thus, we expect that continued supplies of advanced terrestrial prey inputs will keep aquatic prey at a low abundance and litter processing at a suppressed rate.

Another three-month experiment found that prolonged duration of terrestrial prey inputs decreased aquatic prey and litter processing (IX, X, Table 1). This experimental result on aquatic prey may correspond to an instantaneous increase of the model term (2) for the consumption of aquatic prey by small fish. However, our model predictions suggest that long-term effects of prolonged duration are not uniquely determined (Fig. 3). Thus, one possibility is that the empirically observed negative indirect effects of prolonged duration on aquatic prey and litter processing might be halfway toward the eventual reduction of aquatic prey abundance and litter processing rate after multiple years of prolonged terrestrial prey inputs. Yet, another possibility is that the short-term negative indirect effects observed in the experiment could be reversed over a long term, yielding higher aquatic prey abundance and faster litter processing rate. Our model analysis suggests that the long-term effects of prolonged duration depend on total terrestrial prey inputs, pre-perturbed duration, and the growth and reproduction efficiencies of fish. Taking these factors into account will improve long-term predictions for real systems.

Indeed, global warming may not only advance the start of the growing season, but can also extend it (CaraDonna et al. 2014; Post 2018). This might advance and extend the period of terrestrial prey inputs to streams from riparian forests. Our model predicts that advanced timing will generally have a negative effect on aquatic prey and litter processing, but accompanying prolonged duration can either exaggerate the negative effect or have a positive effect to counter the negative effect (Fig. 3B).

In summary, we analyzed the mechanisms by which the timing and duration of terrestrial prey inputs drive indirect effects on aquatic prey and litter processing (Fig. 2). Our analysis clarifies how stage-specific responses of life history processes in fish, such as growth, maturation, and reproduction, mediate and induce indirect effects on the community and ecosystem levels. With growing concerns on potential impacts of climate changes, more research is needed that links how phenological alterations modify species interactions to cause impacts on ecosystem functioning (Donnelly et al. 2011). Stage-specific responses of life history processes are an integral part of the mechanisms with which to predict the consequences of phenological species interactions at the community and ecosystem levels.

## ACKNOWLEDGEMENTS

This work was supported by JSPS Kakenhi 15H04422.

